# Modulation of the host cell transcriptome and epigenome by *Fusobacterium nucleatum*

**DOI:** 10.1101/2021.05.28.445195

**Authors:** Cody A. Despins, Scott D. Brown, Avery V. Robinson, Andrew J. Mungall, Emma Allen-Vercoe, Robert A. Holt

## Abstract

*Fusobacterium nucleatum* (Fn) is a ubiquitous opportunistic pathogen with an emerging role as an oncomicrobe in colorectal and other cancer types. Fn can adhere to and invade host cells in a manner that varies across Fn strains and host cell phenotypes. Here we performed pairwise co-cultures between three Fn strains and two immortalized primary host cell types (colonic epithelial cells and vascular endothelial cells) followed by RNA-seq and ChIP-seq to investigate transcriptional and epigenetic host cell responses. We observed that Fn-induced host cell transcriptional modulation involves strong upregulation of genes related to immune migration and inflammatory processes, such as *TNF, CXCL8, CXCL1*, and *CCL20*. Further, we identified genes strongly upregulated specifically in conditions of host cell invasion, including overexpression of both *EFNA1* and *LIF*, two genes commonly upregulated in colorectal cancer and associated with poor patient outcomes, and *PTGS2* (*COX2*), a gene associated with the protective effect of aspirin in the colorectal cancer setting. Interestingly, we also observed downregulation of numerous histone modification genes upon Fn exposure. To further explore this relationship, we used the ChIP-seq data to annotate chromatin states genome-wide. We found significant chromatin remodeling following Fn exposure in conditions of host cell invasion, with substantial increases in the frequency of states corresponding to active enhancers as well as low signal or quiescent states. Thus, our results highlight increased inflammation and chemokine gene expression as conserved host cell responses to Fn exposure, and extensive host cell epigenomic changes associated with Fn host cell invasion. These results extend our understanding of Fn as an emerging pathogen and highlight the importance of considering strain heterogeneity and host cell phenotypic variation when exploring pathogenic mechanisms of Fn.

## Introduction

*Fusobacterium nucleatum* (Fn) is an elongated rod-shaped, anaerobic, gram-negative bacterium found in both healthy and disease states of human microbiota (Bolstad et al., 1996; Brennan and Garrett, 2019). Fn is ubiquitous in the oral cavity, and can spread beyond its oral niche through the bloodstream, to colonize other body sites (Abed et al., 2016, 2017, 2020; Cochrane et al., 2020). For instance, Fn is an uncommon constituent of the gastrointestinal microbiota in healthy individuals (Amitay et al., 2017; Eklöf et al., 2017).

Fn colonization has been implicated in both oral and extra-oral inflammatory human diseases such as periodontitis (Dzink et al., 1988; Moore and Moore, 1994), endodontic infection (Topcuoglu et al., 2013), inflammatory bowel disease (Strauss et al., 2011), appendicitis (Swidsinski et al., 2011), and adverse pregnancy outcomes (Vander Haar et al., 2018; Han et al., 2010). Moreover, Fn is an emerging oncomicrobe; it is enriched in colorectal cancer (Castellarin et al., 2012; Kostic et al., 2012), breast cancer (Parhi et al., 2020), esophageal cancer (Yamamura et al., 2016) and melanoma (Kalaora et al., 2021). Fn may have a role in tumorigenesis and/or metastatic spread, given it is associated with pre-cancerous colonic polyps (Kostic et al., 2013; McCoy et al., 2013) and with metastatic disease (Bullman et al., 2017; Chen et al., 2020a, 2020b). Clinically, increased Fn tumor burden is associated with poor patient outcomes (Mima et al., 2016), chemoresistance (Yu et al., 2017), and relapse (Serna et al., 2020). Thus, there is a strong rationale to investigate Fn pathogenicity.

It is well established that Fn can adhere to host cells and invade the host cytosol, however, these characteristics vary markedly among Fn strains and host cell types (Han et al., 2000, 2004; Strauss et al., 2011). The virulence mechanisms of Fn and how they may relate to host cell adhesion or invasion are not well understood. The FadA adhesin is a candidate Fn virulence factor, shown to bind E-cadherin and activate Wnt/β-catenin signaling, a pathway frequently dysregulated in colorectal cancers (Bienz and Clevers, 2000; Rubinstein et al., 2013). Fn has also been linked to increased DNA damage (Geng et al., 2020; Okita et al., 2020), possibly related to nuclear localization after invasion (Allen-Vercoe et al., 2011; Han et al., 2004). Further, the invasiveness of Fn strains isolated from inflammatory bowel disease patients correlates with disease severity (Strauss et al., 2011), but it is not known if this is generalizable to other disease settings.

Interaction between Fn and host immunity is similarly complex, with Fn displaying inflammatory and immunogenic properties but also having immunosuppressive effects. For example, Fn induces pro-inflammatory cytokines (Kostic et al., 2013; Park et al., 2014; Rubinstein et al., 2013; Warren et al., 2013) and can stimulate local myeloid cell recruitment (Kostic et al., 2013). Additionally, Fn can also induce adaptive immune responses; colorectal cancer patients have increased levels of anti-Fn antibodies compared to healthy controls (Kurt and Yumuk, 2021; Wang et al., 2016) and anti-Fn CD8^+^ T cells can infiltrate Fn-positive melanoma tumors (Kalaora et al., 2021). However, Fn can also act in an immunosuppressive manner. Fn outer membrane proteins can induce apoptosis in lymphocytes (Jewett et al., 2000; Kaplan et al., 2010) and Fn tumor burden is inversely correlated with CD3^+^ T cell density in colorectal cancer (Mima et al., 2015). Further, Fn inhibits NK and T cell responses via fusobacterial lectin Fap2 binding to the inhibitory TIGIT and CEACAM1 host receptors (Gur et al., 2015, 2019), and Fn can also stimulate tryptophan metabolism to yield kynurenine metabolites that are potent inhibitors of T cell function (Frumento et al., 2002; Xue et al., 2018). Overall, the host-pathogen interactions of Fn are multifactorial and the consequences of Fn tumor enrichment and invasion, such as potential for oncogenic initiation and/or progression, are not fully understood.

Recently, we investigated transcriptional changes in Fn upon host cell invasion and found modulation of genes that may contribute to hematogenous spread and generation of a tumor-permissive hypoxic/inflammatory microenvironment (Cochrane et al., 2020). In the present study, we explore the effects of Fn on the host cell transcriptome and epigenome. We exposed human immortalized primary colonic epithelial and vascular endothelial cells to three strains of Fn *in vitro* using an established invasion assay (Cochrane et al., 2020; Strauss et al., 2011), and then performed RNA-seq and ChIP-seq on the Fn-exposed and unexposed host cell populations. We found that in different conditions Fn exposure upregulated genes related to inflammation (chemokines and cytokines), downregulated genes related to histone modification, and significantly remodeled chromatin states.

## Results

### Fn invades immortalized primary (IP) vascular endothelial cells, but not IP colonic epithelial cells

To investigate the ability of Fn strains to invade different human immortalized primary (IP) cell types, we performed invasion assays using two IP cell lines (human colonic epithelial cells [HCE] and human carotid artery endothelial cells [HCAE]) exposed to three different strains of Fn (subsp. *animalis* 7/1 [Fn 7/1], subsp. *animalis* CC 7/3 JVN3C1 [Fn 7/3], and subsp. *nucleatum* ATCC® 23726™ [Fn 23726]). These colonic epithelial and vascular endothelial host cell lines were chosen to explore consequences of Fn exposure/invasion relevant to colorectal cancer pathogenesis and the capacity of Fn to transit from the bloodstream to other tissues, respectively. After 4 hours of Fn exposure to host cells, invasion efficiency was assessed by microscopy using DAPI nuclear staining and anti-Fn polyclonal antibodies. We found evidence of invasion of HCAE cells by all three Fn strains, and no invasion of HCE cells (**Supplementary Figure 1**).

### Exposure to Fn results in upregulation of host inflammatory responses and downregulation of host histone modification related genes

We repeated the co-incubation of Fn and host cells under each condition in triplicate and extracted host cell RNA for transcriptome analysis (see Methods). We performed differential gene expression analysis independently for each combination of Fn strain and host cell, identifying differentially expressed (DE) transcripts that are upregulated or downregulated in Fn-exposed samples relative to unexposed control cells processed in parallel. For our analysis, we focused on protein-coding genes. Of the 60,671 total annotated human transcripts from Ensembl (Yates et al., 2020), 19,966 are classified as protein-coding.

We first sought to determine the similarity of transcriptional changes for each condition of host cell and Fn strain. We found a large set of significantly DE genes that were shared between all conditions (206 upregulated and 16 downregulated) (**Figure 1A**) (**Supplementary File 1**). We also found hundreds of genes consistently modulated within the same cell line exposed to different Fn strains (host cell-specific changes), while few genes were consistently modulated between the two different cell lines exposed to the same Fn strain (Fn strain-specific changes). This suggests that modulation of the host cell transcriptome by Fn is mainly defined by host cell identity rather than Fn strain identity. Interestingly, however, the HCAE cell line showed an exceptionally large number of uniquely DE genes when exposed to Fn 23726 (828 upregulated and 891 downregulated, compared to 35-202 upregulated and 58-98 downregulated unique genes in other conditions), indicating the potential for context-specific host-pathogen interactions.

**Figure 1.**
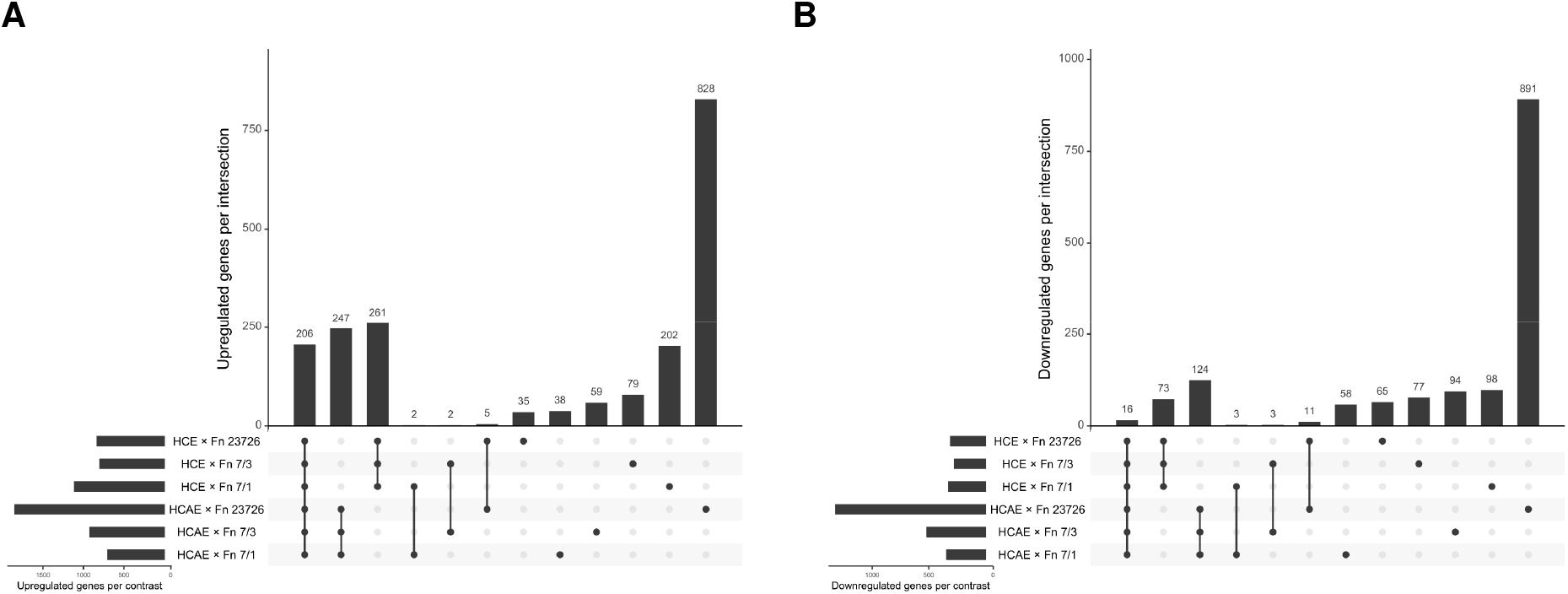
Comparison of HCE and HCAE protein-coding genes differentially expressed in response to various Fn strains. Upset plots highlighting the number of shared significantly differentially expressed upregulated (A) and downregulated (B) genes. Horizontal bars at the bottom left of each plot show the total number of significantly differentially expressed genes in each condition. Vertical bars show the number of genes shared between each comparison, with conditions included in each comparison shown by the respective filled points below.

We next aimed to identify the strongest transcriptional changes in response to Fn exposure in both host cell lines. The mean and standard deviation of each DE gene for each IP cell line exposed to three different Fn strains was calculated (non-significant DE genes log2 fold change (log2FC) = 0). Genes with an average log2FC of > 3 (for upregulated genes) or < −0.75 (for downregulated genes) in both conditions were considered conserved genes of interest (red and blue colored points, top 5 genes of each group labeled) (**Figure 2A**). Using these criteria, we identified 26 genes that were consistently upregulated between conditions of cell lines and Fn strains (*BIRC3, CCL2, CCL20, CSF2, CSF3, CXCL1, CXCL10, CXCL2, CXCL3, CXCL5, CXCL8, ELF3, ICAM1, KCNN3, LTB, NFKBIA, OLR1, OR2I1P, RHCG, RND1, RRAD, TNF, TNFAIP2, TNFAIP3, TNFRSF9, ZC3H12A*). We also identified 17 genes that were consistently downregulated between conditions of cell lines and Fn strains (*AP5S1, ARFGAP2, ASNS, BMI1, ERMAP, FANCE, HRCT1, ID2, KAT6B, KLF15, LPIN2, MAP2K6, MXD3, SH3TC2, TTC30B, ZBTB3, ZNF367*). These are the host genes showing the largest and most consistent expression changes upon Fn exposure.

**Figure 2.**
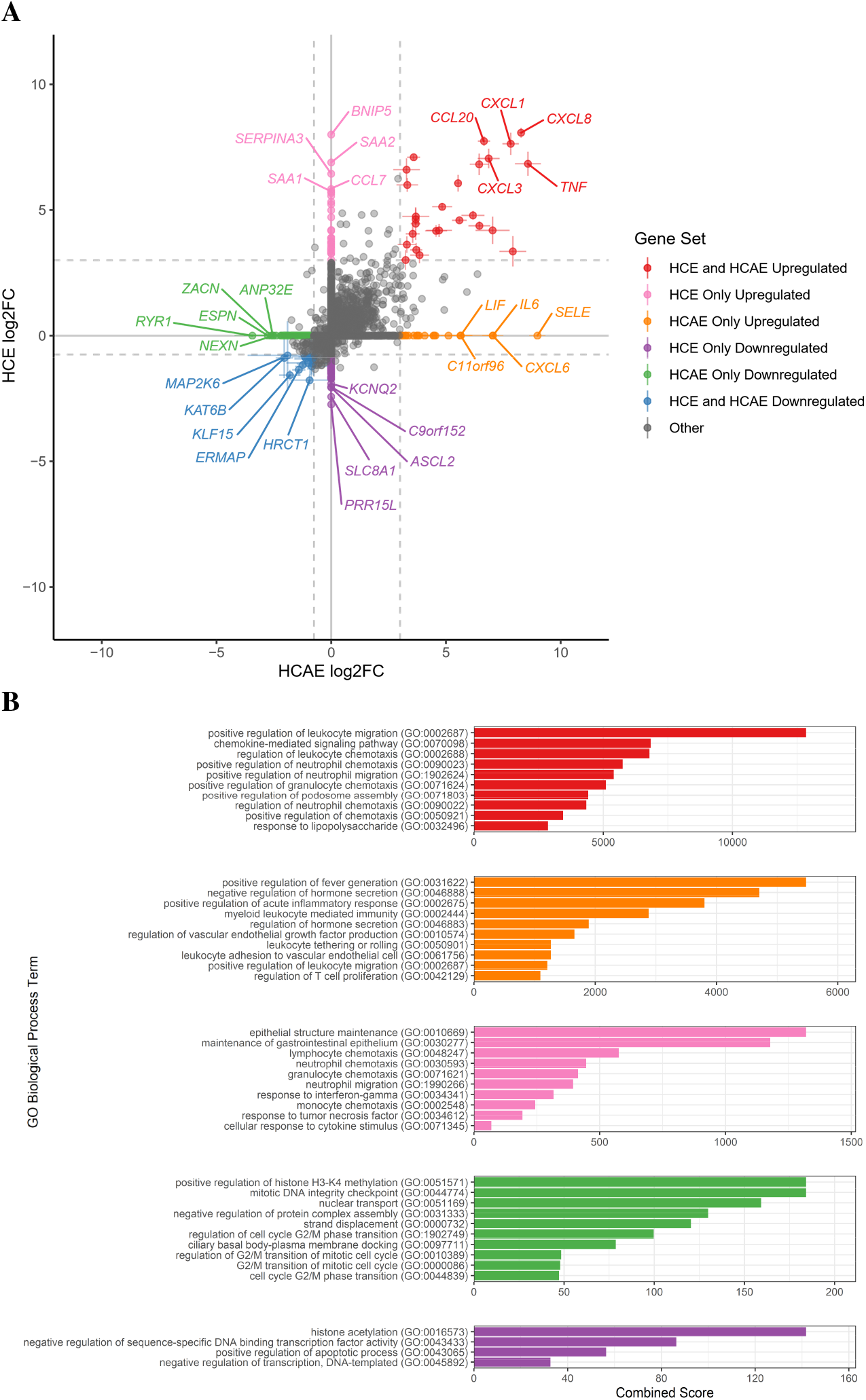
Differential expression gene sets of interest and related pathways. A) Scatterplot and gene set assignments of DE genes for HCE and HCAE. For all genes that were significantly DE (adj. p ≤ 0.05) in at least 1 condition, the mean and standard deviation of DE for each IP cell exposed to three different Fn strains was calculated (non-significant DE genes log2FC = 0, crossbars denote standard deviation). Genes with average log2FC of > 3 (upregulated) or < −0.75 (downregulated) were considered genes of interest (colored points, top 5 genes of each group labeled). Genes meeting these criteria in both cell lines were considered to be conserved upregulated or downregulated genes of interest. Genes meeting these criteria in one cell line and not significantly DE in the other cell line were considered cell line/invasion status specific genes of interest. B) GSEA of gene sets of interest. Ensemble gene names of each gene set were used separately as inputs for EnrichR gene set enrichment analysis (GSEA), querying the GO Biological Processes library. Significant resulting GO terms (adj. p ≤ 0.05) were ranked by combined score (up to 10 terms shown). No significant GO terms were identified for the HCE and HCAE downregulated gene set.

Next, we identified the strongest transcriptional changes in response to Fn exposure that were unique to each host cell line. We considered genes that met our minimum DE cut-offs (log2FC of > 3 or < −0.75) in one cell line and were not significantly differentially expressed in the other cell line as invasion status/cell line specific genes of interest (orange, purple, pink, and green points, top 5 genes of each group labeled) (**Figure 2A**) (**Supplementary File 2**). Given that the HCAE cell line was invaded by Fn and HCE was not, the identified “HCAE only” gene sets represent >250 genes with significant DE only in conditions of invasion in our dataset. While we cannot remove possible confounding effects of host cell genetic background, these genes may include those relevant to invasion-associated Fn pathogenesis.

To explore cellular processes associated with the differentially expressed gene sets of interest, gene set enrichment analysis (GSEA) was performed using EnrichR (Kuleshov et al. 2016). The “GO Biological Process 2018” ontology was queried and significantly enriched terms were ranked by combined score (**Figure 2B**). We observed a number of chemokine and immune cell migration related terms associated with both our conserved upregulated gene set (red), and our cell line-specific upregulated gene sets (orange and pink) (“positive regulation of leukocyte migration”, “chemokine-mediated signaling pathway”, multiple terms relating to chemotaxis of leukocytes/neutrophils/monocytes/granulocytes). Additionally, many terms were related more generally to pro-inflammatory responses (“positive regulation of acute inflammatory response”, “positive regulation of fever generation”, “response to lipopolysaccharide”, “response to interferon-gamma”, “response to tumor necrosis factor”, “cellular response to cytokine stimulus”). This finding suggests that induction of inflammatory and immune recruitment genes dominates the host cell responses to Fn exposure, with additional contributions from cell-type or adhesion/invasion specific differential expression. Interestingly, we also observed cell-line specific terms. For example, using genes only DE in HCE cells as input, the two strongest associated GO terms relate to maintenance of the gastrointestinal epithelium (“epithelial structure maintenance”, “maintenance of gastrointestinal epithelium”. Alternatively, we also noted processes specific to HCAE that align with vascular cell specific functions (“regulation of vascular endothelial growth factor production”, “leukocyte tethering or rolling”, “leukocyte adhesion to vascular endothelial cell”). For GSEA analysis of downregulated gene sets of interest, we found both HCE and HCAE gene lists generated top GO terms relating to histone modification (“histone acetylation” and “positive regulation of histone H3-K4 methylation”, respectively). In addition, a number of terms related to cell cycle progression processes were associated with downregulated genes in HCAE/invaded cells (“mitotic DNA integrity checkpoint”, multiple terms relating to regulation of G2/M phase transition).

We performed GSEA including all significantly upregulated and downregulated genes separately for each combination of host cell and Fn strain to see if any reoccurring terms emerged that may have been hidden by the top gene set analysis. We ranked the significant output GO terms by the sum of their combined scores for each cell line and Fn strain to compare enrichment across all conditions. Similar to trends seen when looking at the strongly upregulated genes of the HCE and HCAE upregulated set, enriched GO terms from the complete set of upregulated genes included terms associated with inflammation and immune cell migration (**Figure 3A**). Emphasizing the importance of results from the conserved downregulated gene subset, a clear trend of histone modification term enrichment emerged, particularly those related to acetylation, when considering all downregulated genes (“histone acetylation” [generally, as well as specific to H3, H4, H4-K5, H4-K8, H4-K12], “positive regulation of histone H3-K4 methylation”, “histone H3-K9 demethylation”) (**Figure 3B**). In this more inclusive analysis, we also observed NF-κB signaling as a previously unidentified top upregulated process (“I-kappaB kinase/NF-kappaB signaling”, “regulation of I-kappaBkinase/NF-kappaB signaling”). Additionally, DNA-damage related processes were associated with downregulated genes (“DNA damage response, signal transduction by p53 class mediator resulting in transcription of p21 class mediator”, “DNA damage response, signal transduction resulting in transcription”).

**Figure 3.**
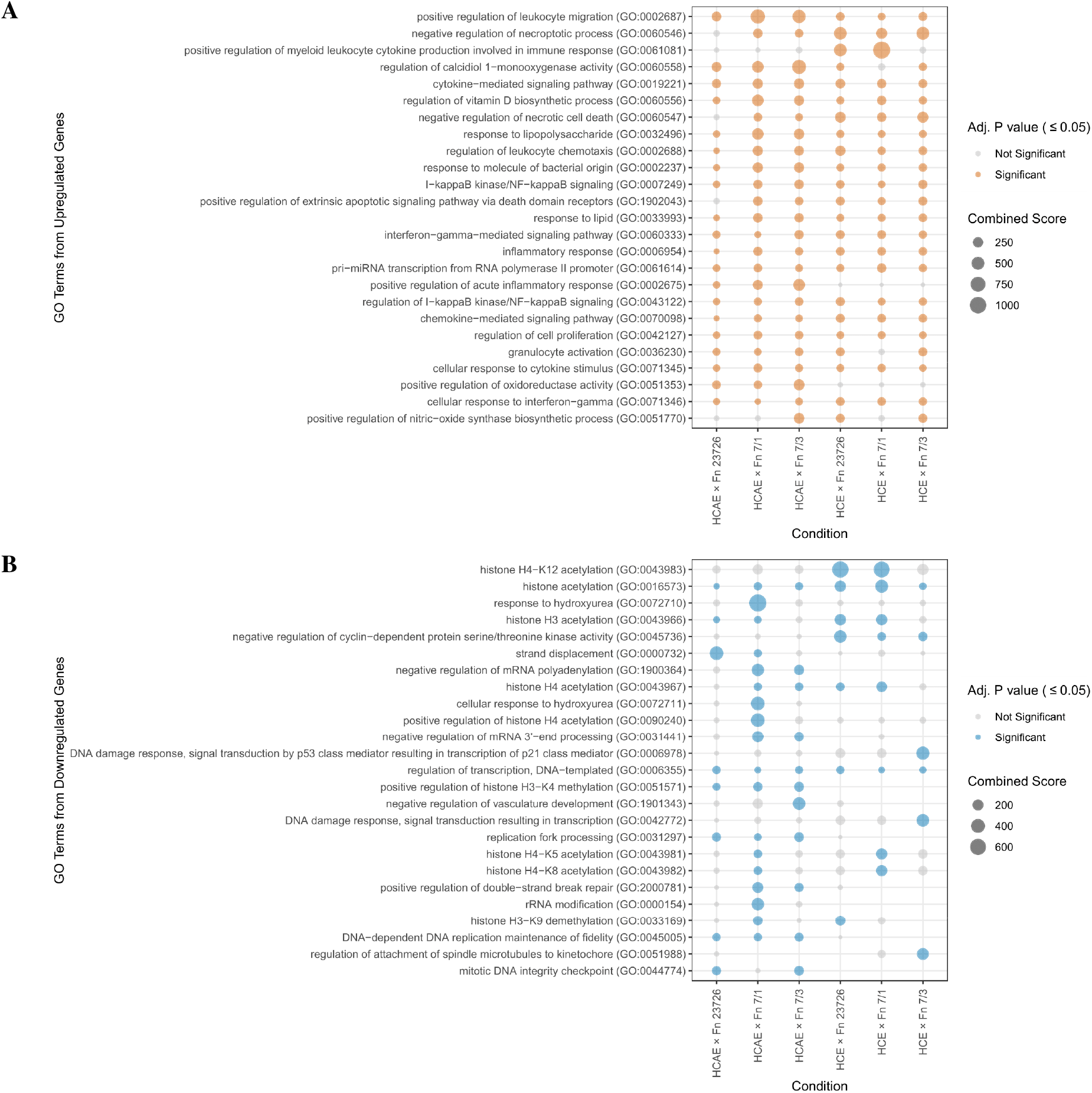
Gene set enrichment analysis (GSEA) of all significantly differentially expressed human protein-coding genes in response to various Fn strains. The y-axis shows the top 25 GO terms for upregulated **(A)** and downregulated **(B)** gene sets for all conditions (ranked by the sum of “Total Score” of significant GO terms from all conditions from top [highest] to bottom [lowest]). Significant GO term total scores (adj. p value ≤ 0.05) are shown as colored (upregulated = orange, downregulated = blue), and non-significant GO term total scores are shown in grey. The size of the dot reflects the score of the GO term for each condition.

### ChIP-seq reveals epigenetic signatures altered by Fn-induced epigenetic changes

Prompted by the observed downregulation of histone methylation/acetylation pathways in the differential gene expression analysis, we evaluated histone modifications in these samples. For these experiments, we exposed HCE and HCAE cells to the Fn 7/1 strain as this well-characterized strain is highly invasive and has been shown to promote tumorigenesis in mice (Kostic *et* al., 2013). We performed ChIP-seq experiments for the six core histone marks used by the International Human Epigenome Consortium (IHEC) (H3K4me1, H3K4me3, H3K9me3, H3K27me3, H3K27ac, and H3K36me3) (Stunnenberg et al., 2016). After alignment of the sequencing data to the human genome, we divided the genome into non-overlapping 200bp windows (n = 15,478,375) using ChromHMM (Ernst and Kellis, 2012), which used the aligned ChIP-seq data to assign the absence or presence of each mark in each of the windows. Next, the overall fraction of the genome containing each mark was compared (**Figure 4A**). In HCAE cells, invadable by Fn 7/1, there was a significant decrease in the fraction of the genome containing H3K4me1 (p = 0.0095, t-test) and a smaller decrease in H3K4me3 (p = 0.019). In HCE, which is not invaded by Fn 7/1, there were no significant changes in the fraction of the genome containing any of the six marks. These observations suggest that Fn invasion influenced the epigenetic state of the HCAE cells.

**Figure 4.**
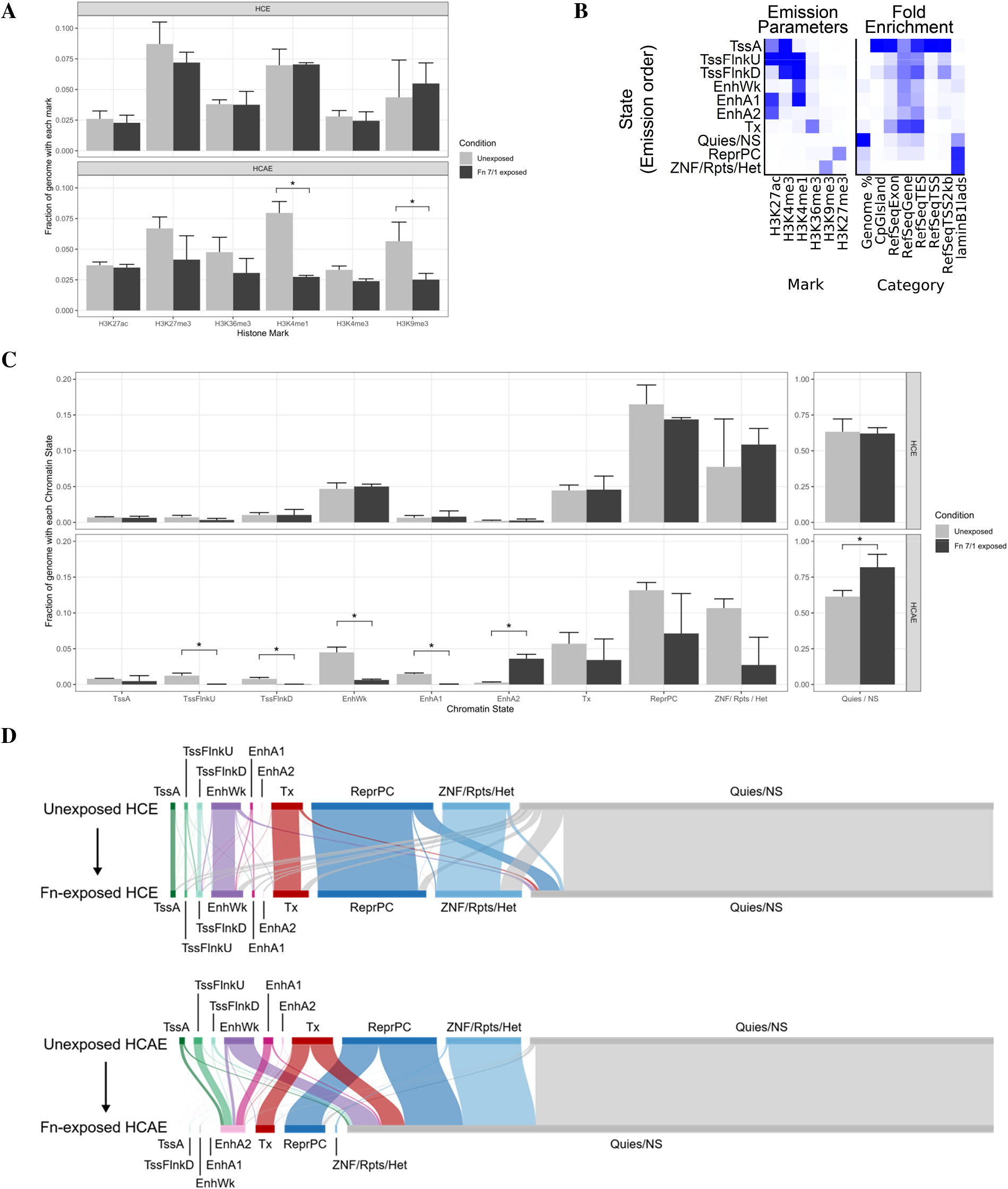
Epigenetic profiling of HCE and HCAE cells exposed to Fn 7/1. **(A)** Barplots showing the fraction of the genome that contains each histone mark (x-axis) in control (light grey) and Fn 7/1 exposed (dark grey) for HCE (top) and HCAE (bottom). Bar height is mean of the three replicates, with the error bar extending to the mean + standard deviation. **(B)** Summary of the 10-state ChromHMM model emission parameters (left panel) and genomic annotations enrichments (right panel). Intensity of blue correlates with increasing values of emission parameter or fold enrichment. **(C)** Barplots showing the fraction of the entire genome that is classified as each state (x-axis) in control (light grey) and Fn 7/1 exposed (dark grey) for HCE (top) and HCAE (bottom). Bar height is mean of the three replicates, with the error bar extending to the mean + standard deviation. The Quies/NS state is shown in its own panel with a different y-axis scale. **(D)** Sankey diagrams showing the changes in states for every window in the unexposed (top) and Fn 7/1 exposed (bottom) for HCE (top diagram) and HCAE (bottom diagram). Bar sizes for each state are proportional to the amount of the genome represented by that state. 96.9 % (HCE) and 96.5 % (HCAE) of the total genome length is represented in these diagrams. Connections from unexposed state (top) to exposed state (bottom) are colored by the unexposed state.

To further explore epigenomic modulation by Fn, we used ChromHMM to create a model of chromatin states. Chromatin states are defined by the combination of histone marks co-occurring in regions of the genome (Ernst and Kellis, 2010; Heintzman et al., 2007; Wang et al., 2008). We selected a model with ten chromatin states based on published chromatin state models (Gorkin et al., 2020; Roadmap Epigenomics Consortium et al., 2015) and manual inspection of the emission parameters and genomic characteristics of each state (**Figure 4B**). Our ten chromatin states can be divided into five groups; Promoter (Active TSS [TssA], Flanking Active TSS Upstream [TssFlnkU], Flanking Active TSS Downstream [TssFlnkD]), Enhancers (Weak Enhancer [EnhWk], Active Enhancer 1 [EnhA1], Active Enhancer 2 [EnhA2]), Genic Transcription (Transcription [Tx]), Repressed (Repressed Polycomb [ReprPC], ZNF Genes, Repeats, Heterochromatin [ZNF/Rpts/Het]), and Low Signal (Quiescent / No Signal [Quies/NS]). Using this model, we assigned every 200 bp window of the genome its best-evidenced chromatin state. When looking at the fraction of the genome that was classified as being in each state, we observed that for both HCE and HCAE, the majority of the genome was classified as the Quies/NS state (**Figure 4C**).

Changes in the genome-wide abundance of chromatin states in response to Fn exposure were strikingly different in the invaded HCAE cells compared to the non-invaded HCE. In HCAE we saw significant decreases in TssFlnkU, TssFlnkD, EnhWk, and EnhA1, and increases in EnhA2 and Quies/NS (p < 0.05). The largest changes in HCAE were in the “Quies/NS” state (20.6% increase) and “EnhWk” (3.9% decrease). In HCE there are no significant changes in any of the states. We summarized the specific state changes undergone by each window of the genome in a Sankey diagram (**Figure 4D**). This view highlights the predominant state transitions in HCAE cells upon Fn exposure, including substantial shifts from the TssFlnkU and EnhA1 states to the EnhA2 state, and from the EnhWk, Tx, ReprPC and ZNF/Rpts/Het states to the Quies/NS state.

### Fn-associated chromatin state changes correlate with Fn-associated gene expression changes

Given that we found regions of the HCAE genome with changes in chromatin state upon exposure to Fn, we sought to correlate the specific state changes with changes in gene expression. We identified 4,456,888 windows in the HCAE genome that changed upon Fn exposure. To ascribe the epigenetic impact of these windows to specific genes, we used GREAT (McLean et al., 2010) to associate each genomic region with one or more genes using the basal regulatory region rules (see Methods). To assign a quantitative value to the cumulative effect on transcription for genes with associated state changes, we scored each state change based on the predicted effect of the state on gene expression (+2: TssA, Tx. +1: TssFlnkU, TssFlnkD, EnhWk, EnhA1, EnhA2. 0: Quies/NS. −1: ReprPC, ZNF/Rpts/Het) and determined the difference in value between the ending state and starting state to assign an “epigenomic score” value to each window. Then, we summed all epigenetic scores for all windows associated with a specific gene to get a Net Epigenomic Score for each gene. Focusing on each set of genes identified in Figure 2A, we measured the correlation between the log2FC and Net Epigenomic Score for these genes and tested for significance by a random resampling method (see Methods). We saw significant correlations for HCAE upregulated genes (Fig 2A, HCE and HCAE upregulated [red points], r = 0.559, p = 0.02; HCAE only upregulated [orange points] r = 0.506, p = 0.03) and downregulated genes (Fig 2A, HCE and HCAE downregulated [blue points]; r = 0.517, p = 0.04). Repeating this analysis for HCE cells, we saw no significant correlations between expression changes and epigenomic changes, further substantiating the sensitivity of the invadable HCAE cell line to epigenomic modulation by Fn.

## Discussion

*Fusobacterium nucleatum* (Fn) is among the most common bacterial species of the oral cavity and is becoming increasingly recognized as an opportunistic pathogen, implicated in diseases including numerous cancer types and both oral and extraoral infections. Despite such inculpatory findings, understanding of Fn pathogenesis and virulence factors that contribute to disease development is complicated by Fn strain diversity and differential effects in host cells and ultimately remains limited.

Here, we have comprehensively profiled transcriptional and epigenetic host cell changes in response to Fn exposure. To address possible differences in host cell response between Fn strains and host cell types, we performed pairwise invasion assays for 2 different human immortalized primary cell lines using 3 separate Fn strains.

Our results indicate that Fn exposure upregulates genes related to immune migration and inflammatory processes, consistent with previous reports. For example, Fn treatment increased tumor-infiltrating myeloid cell density and chemokine expression in the Apc^*min*/+^ mouse model (Kostic et al., 2013). The human colorectal cancer derived HCT116 cell line has been shown to upregulate *CXCL8* (*IL-8*) and *CXCL1* in response to Fn 23726 and may contribute to metastatic spread (Casasanta et al., 2020). Modulation of these genes is corroborated in our data, with *CXCL8* and *CXCL1* among the highest upregulated genes in both HCE and HCAE cell lines upon exposure to all Fn strains. *TNF* has also been reported to be induced by multiple invasive Fn strains in LS 174T colonic cells (Dharmani et al., 2011). We show that this pro-inflammatory cytokine is upregulated strongly by both HCE and HCAE cells. *CCL20* recruits Treg cells and is associated with chemoresistance in the setting of colorectal cancer (Wang et al., 2019). Upregulation of *CCL20* and *CSF-3* (*G-CSF*) has been shown in Fn-infected esophageal tumor tissue and gingival fibroblasts, respectively (Kang et al., 2019a; Yamamura et al., 2016). *CSF-2* (*GM-CSF*) upregulation has been observed in Fn-infected gingiva-derived mesenchymal stem cells (Kang et al., 2019b), and a role for *CSF-2* in CRC epithelial-mesenchymal transition has been proposed (Chen et al., 2017). We show upregulation of *CCL20, CSF-3*, and *CSF-2* is induced in both HCE and HCAE cells upon exposure to all Fn strains tested in our study. These collective findings therefore suggest that upregulation of these genes may be an important feature of Fn-induced host responses consistent between various host cell settings and Fn strains, and our data indicate this modulation is not strictly dependent upon Fn host cell invasion. Overall, our results extend the emerging trend of inflammatory and immune migration related gene upregulation as a conserved key aspect of host response to Fn exposure.

Gene set enrichment analysis (GSEA) also revealed several notable pathways modulated upon Fn exposure. Fn has been previously shown to increase host cells in S phase and G2/M phase (Ma et al., 2018). We show G2M cell cycle related terms are among the most enriched Biological Process GO terms associated with the HCAE specific downregulated gene set. The genes related to these terms (*AKAP9, CDK5RAP2, CEP57, CEP135, CEP290, CNTRL, KIF14, NINL, PPP1R12A, USP47*) therefore may contribute to changes in host cell proliferation following exposure to Fn in some host cell contexts. Additionally, we observed upregulation of genes related to gastrointestinal maintenance in HCE cells (*MUC2, SERPINA3*), possibly in response to disruption of gastrointestinal barrier integrity by Fn (Liu et al., 2020).

Our analysis also highlights genes that may be differentially expressed by host cells in response to Fn invasion. *PTGS2* (*COX2*) upregulation is associated with Fn abundance in colon tumors (Kostic et al., 2013). Additionally, aspirin, a modulator of *PTGS2*, has been recently demonstrated to reduce development of colonic adenomas after Fn inoculation in the Apc^*min*/+^ mouse model (Brennan et al., 2021). In our data, *PTGS2* is among the most strongly differentially expressed genes in invaded HCAE cells, but is not differentially expressed in HCE cells, suggesting that upregulation of *PTGS2* and aspirin inhibition may be particularly important for invasive Fn strains. Additionally, we found *EFNA1* (Ephrin A1) and *LIF* (leukemia inhibitory factor) are also differentially expressed only in conditions of invasion. Overexpression of both *EFNA1* and *LIF* is common in colorectal cancer and associated with poor patient outcomes, and *LIF* expression mediates chemoresistance via negative regulation of p53 *in vitro* (Wu et al., 2015; Yamamoto et al., 2013). Thus, these genes and others observed to be modulated only in conditions of invasion in this study represent invasion-specific candidates that can be evaluated for their relationship to pathogenesis and correlation with disease severity in future studies.

Previously, Hong et al. (2020) reported histone acetylation of specific genes following Fn-induced lncRNA ENO1-IT upregulation. To our knowledge, our present study is the first to explore epigenome-wide changes in host chromatin in response to Fn exposure. Strikingly, we saw significant epigenomic changes in HCAE cells but not in HCE cells, suggesting that these changes may be linked to host cell invasion. Indeed, it has been observed that following invasion Fn can localize to the nucleus (Allen-Vercoe et al., 2011; Han et al., 2004). Further, we showed that the level of differential gene expression in highly differentially expressed genes correlates with these invasion-related chromatin changes, supporting invasion-induced epigenomic restructuring and transcriptomic dysregulation.

It is interesting to note that when looking at the transcriptome, HCE and HCAE cells have similar overall responses to Fn exposure, with a strong upregulation of inflammation-related genes, however epigenomic changes are largely restricted to HCAE cells. One model that could explain this observation is that Fn exposure leads to an initial host cell transcriptional response in both HCE and HCAE cells. Then, in HCAE cells that Fn can invade, intracellular Fn influences changes to histone marks, leading to changes in chromatin states and further modulating gene expression.

Our results should be interpreted within the context of our experimental design. As it is not possible to directly compare the effects from invasion versus exposure alone for a given Fn strain × host cell pair, we are unable to definitively discriminate between effects related to invasion and exposure in invaded HCAE cells. Additionally, we recognize that the sensitivity to detect invasion-related DE is limited by a proportion of host cells remaining un-invaded by Fn in co-culture. The co-cultures are time-limited because the aerobic conditions needed by host cells are unfavorable to anaerobic bacteria like Fn. In future studies, DE specific to invaded cells can be explored using single-cell sequencing methods.

In summary, we have identified candidate genes and gene pathways that are dysregulated in host cells co-cultured with Fn. Additionally, we have shown genome-wide and specific chromatin state changes associated with Fn invasion which correlate with observed changes in gene expression. Our results highlight upregulation of inflammation and chemokine gene expression and downregulation of histone modification related genes as common host cell responses to Fn exposure and demonstrate epigenetics as a new area of research towards defining Fn pathogenicity.

## Materials and Methods

### Immortalized primary cell maintenance and preparation for co-incubation

Two immortalized primary (IP) cell lines were obtained from suppliers: human carotid artery endothelial cells (HCAE) [ABM T0512] and human colonic epithelial cells (HCE) [ABM T0570]. HCAE and HCE cells were maintained in PriGrow I and III (ABM), respectively, supplemented with 10% FBS and 50 μg/mL plasmocin. IP cells were allowed to reach 100% confluence before washing with 0.2 μm filter-sterilized phosphate-buffered saline (PBS) and were harvested using 0.25% trypsin-EDTA (Thermo Fisher) for 5 mins at 37°C, 5% v/v CO_2_. Following trypsin quenching by addition of respective cell medium, IP cells were collected by centrifugation at 300 x *g* for 3 min and seeded into a new flask at a 1:5 seeding ratio. Once IP cells reached 70% confluence at passages 2-7, the media was aspirated, and the cells were washed gently in PBS. Serum- and antibiotic-free DMEM was then applied in preparation for co-incubation.

### Fn strain maintenance and preparation for co-incubation

Three Fn strains were used in this study: subsp. *animalis* 7/1 (Fn 7/1, also reported as EAVG_002) isolated from an inflammatory bowel disease patient (Strauss *et al*., 2011), subsp. *animalis* CC 7/3 JVN3C1 (Fn 7/3) isolated from a colorectal cancer patient (Cochrane *et al*., 2020), and subsp. *nucleatum* ATCC 23726 (Fn 23726). Fn strains were maintained in a humidified anaerobic chamber (Ruskinn Bug Box) at 37°C in 10% v/v CO_2_, 1% v/v H_2_, balance N_2_ on Fastidious Anaerobe Agar (Neogen) supplemented with 5% v/v defibrinated sheep’s blood (Hemostat Laboratories) (FAA). Fn broth cultures for co-incubations were grown overnight (16-18 hr) to late-log phase in TSB supplemented with 5 μg/mL hemin and 1 μg/mL menadione, 0.2 μm filter-sterilized (supp. TSB) that had been previously incubated anaerobically for >16 hr (degassed). The Fn cultures were then diluted with degassed supp. TSB to 10^8^ c.f.u. ml^−1^, verified by MacFarland standards.

### Co-incubation of immortalized primary cells and Fn

Co-incubations between Fn and human cells were completed in biological triplicate at separate human cell passages. Diluted Fn broth cultures were applied to IP cells at a MOI of 50:1 fusobacterial cells:host cells. IP cells were incubated with Fn 7/1, or Fn 7/3, or Fn 23726, or alone (control) for 4 hr at 37°C, 5% v/v CO_2_.

### Immunofluorescent visualization of Fn and immortalized primary cell co-incubations

IP cells were grown to 70% confluence at passage 2 on coverslips pre-treated with poly-L-lysine (Sigma). Briefly, coverslips were incubated for 5 mins in 1 mg/mL poly-L-lysine at room temperature then washed once with sterile deionized water. Coverslips were allowed to dry >2 hr at room temperature before seeding with IP cells at 1:5 seeding ratio. Co-incubations of each Fn strain and IP cell line were completed in biological triplicate. Following 4 hr co-incubation, cells were gently washed twice with PBS. Cells were then fixed with 4% paraformaldehyde (Sigma) for 15 mins at room temperature. Paraformaldehyde was then aspirated and cells were blocked >24 hr with 10% normal goat serum (NGS) (Thermo Fisher) at 4°C. External Fn cells were stained for 30 min at 37°C with primary rat anti-Fn antibodies: Fn 7/1 with EAV_AS4 (diluted 1/500 in NGS), Fn 7/3 with EAV_AS2 (diluted 1/200 in NGS), and Fn 23726 with EAV_AS7 (diluted 1/500 in NGS) (Strauss et al., 2011). EAV_AS7 was prepared alongside previously published rat antibody sera and has been determined to be reactive with Fn 23726 by immunofluorescence microscopy (data not shown). Cells were then washed vigorously for 5 seconds three times with PBS. Goat anti-rat IgG antibodies conjugated to Cyanine3 (Cy3) (Thermo Fisher) diluted 1/500 in NGS containing 1 µg/mL DAPI were then applied to the cells. Cells were allowed to incubate for 30 mins at room temperature, in the dark. Cells were then washed vigorously for 5 seconds three times with PBS. Coverslips were mounted on microscopy slides by applying MOWIOL 4-88 (Sigma) and allowing to solidify overnight (14-20 hr), in the dark. Fluorescent micrographs of stained IP cells and Fn cells were collected on a Leica DM 5000B using Semrock – DAPI and Texas Red filters.

### Cell harvest for RNA extraction

Following 4 hr co-incubation, cells were gently washed twice with PBS. Cells were harvested using 0.25% trypsin-EDTA (Thermo Fisher) for 5 mins at 37°C, 5% v/v CO_2_, neutralized with the respective medium containing 10% v/v FBS, and collected by centrifugation at 300 x *g* for 3 min. Cells were then resuspended in 1 mL TRIZol (Thermo Fisher), incubated for 3 min at room temperature, and then frozen at −20°C until RNA harvest. RNA was extracted from the TRIZol samples according to manufacturer standards. Briefly, samples were allowed to come to room temperature then were combined with 0.2 mL chloroform. Samples were incubated 2 minutes prior to centrifugation at 12,000 x *g* for 15 min at 4°C. The aqueous phase was carefully removed by pipetting and combined with 0.5 mL isopropanol. The mixture was incubated 10 min, then centrifuged at 12,000 x *g* for 10 min at 4°C. The supernatant was carefully discarded by pipetting. The RNA pellet was then washed twice in 75% ethanol, collected by centrifuging at 7,500 x *g* for 5 min at 4°C, and then air-dried for 10 min. The RNA pellet was then resuspended in 50 µL nuclease-free water and incubated 10 min at 55°C. The RNA was then treated with dsDNase (Thermo Fisher) according to manufacturer standards. The quantity and quality of the RNA was assessed by Agilent TapeStation 4150 analysis. All samples submitted for RNA sequencing had RIN values > 7.5.

### Cell harvest for chromatin immunoprecipitation

Following 4 hr co-incubation of Fn 7/1 with each of the IP cells or IP cells alone, cells were gently washed twice with PBS and harvested using 0.25% trypsin-EDTA at 37°C, 5% v/v CO_2_. Trypsin was then neutralized with the respective medium containing 10% v/v FBS, and cells were collected by centrifugation at 300 x *g* for 3 min. Cells were then washed once in PBS and harvested by centrifugation at 300 x *g* for 3 min. PBS was then aspirated and cells were immediately snap-frozen in liquid N_2_. Frozen cell samples were stored at −80°C prior to analysis.

### Chromatin extraction and immunoprecipitation

Generation of ChIP-seq libraries from frozen cell pellets was performed by Canada’s Michael Smith Genome Sciences Centre, Vancouver, Canada. For each sample, immunoprecipitation was performed for six core histone marks (H3K9me3, H3K27me3, H3K36me3, H3K4me1, H3K4me3, and H3K27ac). ChIP libraries were made for each of these marks, and one for input control DNA, yielding seven libraries per sample. Libraries were constructed with IDT’s Duplex UMIs for improved duplicate read detection.

### RNA sequencing

RNA sequencing was performed by Canada’s Michael Smith Genome Sciences Centre, Vancouver, Canada. For each sample, ribo-depleted strand-specific RNA-seq libraries were constructed. All sequencing was performed using 150bp paired-end reads on an Illumina HiSeq 2500 instrument. Each of the 24 libraries was individually barcoded, pooled, and sequenced at approximately 10 libraries per lane. Results of sequencing are summarized in Supplementary File 3.

### ChIP sequencing

ChIP sequencing was performed by Canada’s Michael Smith Genome Sciences Centre, Vancouver, Canada. Libraries were sequenced using 150bp paired end reads on an Illumina HiSeqX, at a target depth of 100 million reads for H3K9me3, H3K27me3, H3K36me3, H3K4me1, and Input Control DNA, while H3K4me3 and H3K27ac had a target depth of 50 million reads. Results of sequencing are summarized in Supplementary File 3.

### RNA-seq data processing

RSEM (Li and Dewey, 2011) `rsem-calculate-expression` was used to align raw RNA-seq datasets to the GRCh38 reference genome and calculate gene expression values (ensembl101) (Yates et al., 2020). Differential gene expression was assessed using DESeq2 (Love et al., 2014), using the established workflow for processing RSEM-generated data (setting the minimum gene length to 1 for all genes after running `tximport()` and before running `DESeqDataSetFromTximport()`). This workflow includes estimation of size factors, estimation of dispersion, and negative binomial GLM fitting. Log fold change shrinkage was performed using the Normal prior.

### RNA-seq data analysis

Differentially expressed genes were selected by an adjusted p-value threshold of 0.05. Ensembl IDs were mapped to gene names using ensembl101. To define top gene sets of interest associated with various conditions, the mean and standard deviation of DE for each IP cell exposed to three different Fn strains was calculated for all genes that were significantly DE (adj. p ≤ 0.05) in at least 1 condition, (non-significant DE genes log2FC = 0). Genes with average log2FC of >3 (upregulated) or < −0.75 (downregulated) were considered genes of interest for the respective condition(s), as shown in Figure 2A.

### GSEA of differentially expressed genes

Top gene sets of interest shown in Figure 2A, as well as all significantly differentially expressed genes (selected by an adjusted p-value threshold of 0.05), had ensembl IDs mapped to gene names using ensembl101. These gene names were used as input for enrichR. In Figure 2B, GO terms were ranked by the “Total Score” of significant GO terms from top (highest) to bottom (lowest). In Figure 3, GO terms were ranked by the sum of “Total Score” of significant GO terms from all conditions from top (highest) to bottom (lowest).

### ChIP-seq data processing

UMIs were extracted and sequences were mapped to human reference hg19 via BWA-MEM. Deduplication was performed with picard tools MarkDuplicates. ChromHMM was used to binarize the aligned ChIP-seq data for each replicate individually. In the ChromHMM design file, the Input DNA ChIP-seq data was provided as a local control for each mark to control for local variances in genome coverage. Chromatin state models containing 2-24 states were learned using the LearnModel command, and all models were compared to the Roadmap Epigenomics Consortium (REC) (Roadmap Epigenomics Consortium et al., 2015) 18-state model which was defined using the same six histone marks we investigated. By assessing the median correlation of the 18 states in each of our potential models, we found 10 states to be the smallest number of states which maximized the correlation. By correlating the emission parameters of each state in our 10-state model to the emission parameters of the REC 18-state model, and by manual inspection of the genomic annotations, we classified our states based on the REC 18-state model.

### ChIP-seq data analysis

To obtain a higher confidence set of chromatin state annotations for our samples, we identified the regions which had consistent states called across each of the three replicates. Here, we defined a consistent state as at least two of the three replicates with the same state assignment, and we scanned through the genome to identify each consistent 200bp window. Windows which did not have matching states across at least two of the three replicates in both the unexposed and Fn exposed samples were categorized as not having a consistent state and were removed from further analysis. Having a single state call for every 200bp window for each sample, we then compared all regions in the control and Fn exposed samples to identify the state change associated with each region which correlates with Fn exposure.

### Integration of transcriptomic and epigenomic data

To integrate the chromatin state changes with transcriptomic changes, we first used GREAT (McLean et al., 2010) to find gene associations for these regions using the default “Basal plus extension” association rule modified to not allow Distal associations (0 kb) and not including curated regulatory domains. We then defined a Net Epigenomic Score for each gene which quantifies the summative effect of all associated windows on the gene. States were assigned scores based on the predicted effect on transcription (+2: TssA, Tx. +1: TssFlnkU, TssFlnkD, EnhWk, EnhA1, EnhA2. 0: Quies/NS. −1: ReprPC, ZNF/Rpts/Het), and the score for each window was calculated as the score for the state in the Fn exposed sample subtract the score for the state in the unexposed sample. The Net Epigenomic Score was calculated for each gene by taking the sum of scores for all windows associated with that gene. To test the correlation with gene expression changes, we calculated the Pearson correlation coefficient between the gene expression (log2FC) and Net Epigenomic Score for subsets of genes identified in the gene expression analysis. To test the statistical significance of these correlations, we randomly sampled equally sized sets of genes and tested the correlation for these random subsets, repeated this 1000 times, and recorded the p-value as the fraction of these 1000 trials which resulted in a correlation coefficient more extreme (farther from zero) than the observed correlation coefficient.

## End Matter

### Data and Code Availability

The RNA-seq and ChIP-seq data generated in this study have been submitted to the NCBI BioProject database (https://www.ncbi.nlm.nih.gov/bioproject/730807) under accession number PRJNA730807. Custom code used in this study is available in the following GitHub repository: https://github.com/scottdbrown/host-transcriptome-epigenome-fuso.

## Author Contributions

R.A.H and E.A.V designed research, A.V.R. performed invasion assays, A.J.M. coordinated sequence data generation, C.A.D. and S.D.B analyzed data, and C.A.D, S.D.B. and R.A.H. wrote the paper.

Author E.A.V. is co-founder and CSO of NuBiyota LLC, a company that is developing microbial ecosystem therapeutic drugs for use in a variety of human disease indications.

## Acknowledgments

This research was funded by the Canadian Cancer Society grant to R.A.H. and E.A.V. (grant #705314). C.A.D. and A.V.R were supported by Canadian Graduate Scholarships from the Canadian Institutes of Health Research (CIHR). We would like to thank the staff of Canada’s Michael Smith Genome Sciences Centre at BC Cancer for performing library construction, sequencing, and data QC. We would also like to thank Dr. Michaela Strüder-Kypke at the Molecular and Cellular Imaging Facility, Advanced Analysis Centre, University of Guelph, for her assistance with confocal fluorescence microscopy.

## Supplementary Data

**Supplementary Figure 1.**
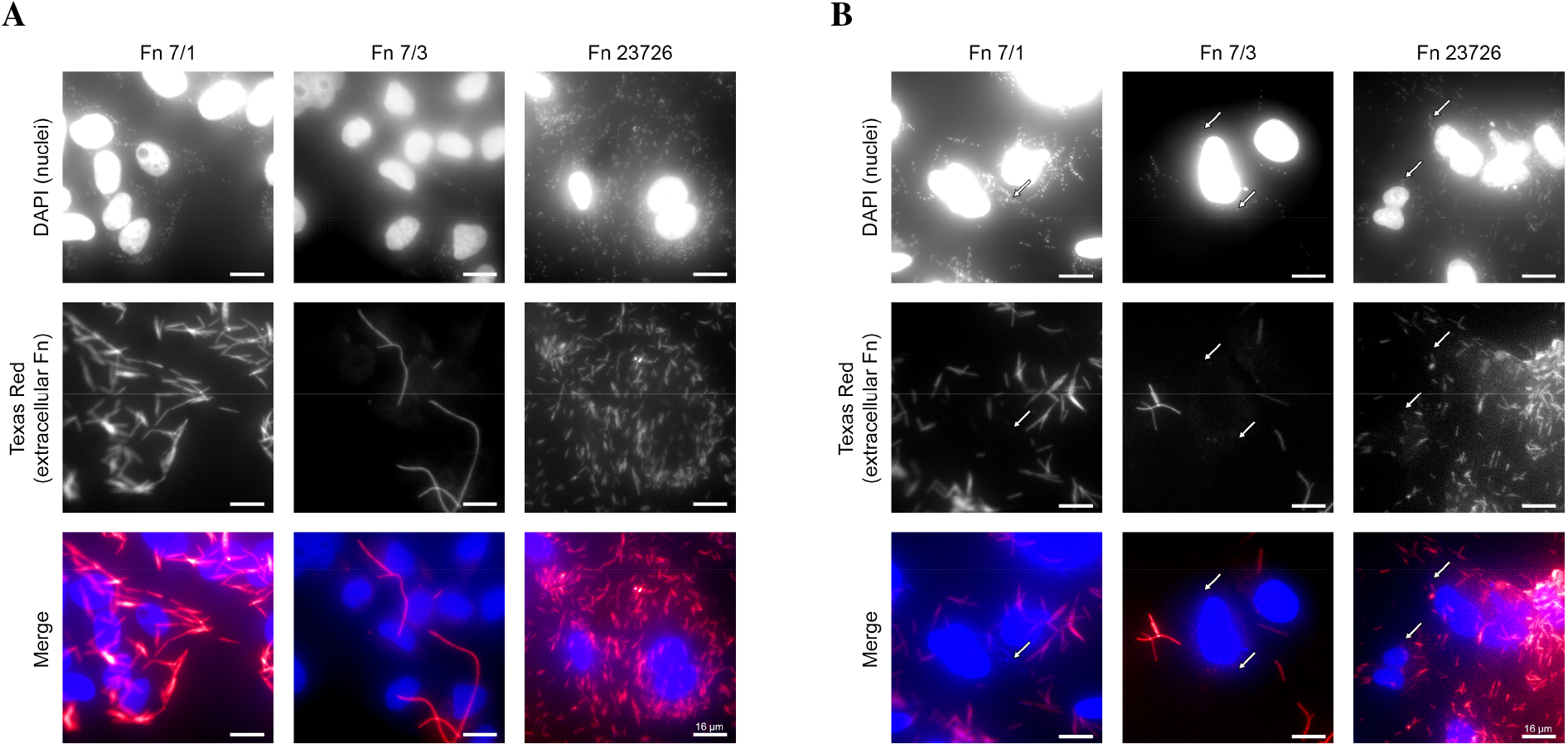
Representative fluorescent microscopy images of each host cell × Fn strain condition. HCE (A) and HCAE (B) host cell lines were stained with DAPI, primary rat anti-Fn antibodies, and secondary goat anti-rat IgG antibodies conjugated to Cyanine3 (Cy3). DAPI (blue) stains host cell nuclei as well as Fn cell nuclei of intracellular and extracellular Fn. Cy3 Fn staining (red) only labels extracellular Fn. Fn nuclei appear as punctate blue points along the elongated Fn body. White arrows highlight invaded Fn, characterized by blue points near the host cell nuclei without any red staining.

## Notes

https://www.ncbi.nlm.nih.gov/bioproject/730807

https://github.com/scottdbrown/host-transcriptome-epigenome-fuso

